# Nod factor signaling controlled genes in *Medicago truncatula* nodules

**DOI:** 10.64898/2026.06.23.733963

**Authors:** Jieyu Liu, Defeng Shen, Erik Limpens, Elena E. Fedorova, Rik Huisman, Huchen Li, Tian Zeng, Joël Klein, Hetty C. van den Broeck, Elio Schijlen, Olga Kulikova, Ton Bisseling

## Abstract

- Legume nodule formation is induced by rhizobia secreted Nod factors (NFs). It has been shown that NF receptors also accumulate in the apex of *Medicago truncatula* nodules. However, the NF signaling induced transcriptional changes in there have never been studied.
- Here, we studied this by using NF signaling mutant TE7, a weak allele of *IPD3,* blocked in rhizobial release. Nodule apices were isolated with laser microdissection and used for transcriptional analysis.
- We identified 1655 NF signaling controlled genes in nodule apex. By comparing this with the transcriptome data from *VAMP721d&e RNAi* nodule apices, we identified a subset of 445 genes whose expression depends on NF signaling and rhizobial release. Further, we compared the set of genes controlled by NF signaling in nodule apices with that controlled in root epidermis, and these showed only a small overlap. *NIN* is induced by NF signaling both in the root epidermis and in the nodule. By overexpression of *NIN* in TE7 and knock down of *NIN* in wildtype nodules we showed that NF signaling controlled rhizobial release depends on NIN.
- NF signaling controls a distinct set of genes in nodules, the function of which depends at least in part on NIN.

## Introduction

By establishing a root nodule symbiosis with nitrogen-fixing rhizobia, legumes can grow in nitrogen-poor soil. In most legume species, the formation of these nodules is induced by lipo-chito-oligosaccharides, known as Nod factors (NFs), that are secreted by the rhizobia (Denarie and Debelle, 1996). NFs are perceived in the root epidermis by LysM domain containing receptor like kinases, which activate a specific signal transduction cascade (Downie, 2014). This leads to rapid transcriptional changes in the root and sets in motion the nodule formation process. In mature nodules, numerous genes are specifically upregulated in comparison to roots. However, how rhizobia induce these genes remains largely unknown. In *Medicago truncatula* (Medicago) it has been shown that NF receptors accumulate at the apex of mature nodules (Moling et al., 2014). We hypothesize that NF signaling contributes to transcriptional changes in the nodule apex leading to the formation of nitrogen fixing organelles, the symbiosomes. In this study, we aimed to identify genes that are regulated by NF signaling in the Medicago nodule apex.

NFs are perceived by membrane located heteromeric complexes of LysM-domain receptor kinases (LYK3/NFP in Medicago; NFR1/NFR5 in *Lotus japonicus* [Lotus]) (Amor et al., 2003; Limpens et al., 2003; Madsen et al., 2003; Radutoiu et al., 2003). After perception of NFs at the root epidermis, a signal transduction pathway is activated. This pathway involves several components, including a leucine-rich repeat receptor kinase (DMI2 in Medicago and SYMRK in Lotus) (Endre et al., 2002; Stracke et al., 2002), nuclear membrane-located cation channels (DMI1 in Medicago; CASTOR and POLLUX in Lotus), a calcium calmodulin-dependent kinase (DMI3 in Medicago and CCaMK in Lotus) and transcription factors (IPD3 and IPD3-like in Medicago; CYCLOPS in Lotus) (Mitra et al., 2004; Tirichine et al., 2006; Messinese et al., 2007; Yano et al., 2008; Jin et al., 2018). Perception of NFs leads to the activation of CCaMK (Mitra et al., 2004; Tirichine et al., 2006), which subsequently activates the transcription factor CYCLOPS by phosphorylation (Messinese et al., 2007; Yano et al., 2008). Subsequently, CYCLOPS induces the expression of *NIN* (*NODULE INCEPTION*). NIN is a transcription factor that is essential for >95% of all transcriptional changes induced in the roots 24h after inoculation with rhizobia, underlining its importance in NF induced transcriptional changes in roots (Schiessl et al., 2019).

NF signaling activates two main processes during nodule initiation; the infection process, that starts in the epidermis, and the mitotic activation of inner root cells, resulting in the formation of a nodule primordium (Oldroyd and Downie, 2008; Xiao et al., 2014). NIN is essential for both processes (Schauser et al., 1999; Marsh et al., 2007). Infection often involves the formation of a tube-like structure, named infection thread, by which the rhizobia enter the plant. Infection threads grow towards the primordia and enter the mitotically activated cortical cells. Subsequently, rhizobia are released from the infection thread and a nodule is formed (Xiao et al., 2014).

Medicago forms indeterminate nodules, which have a persistent apical meristem that adds cells to the nodule tissues (Hirsch, 1992; Sprent, 2007). These are the central tissue, containing the infected cells that are interspersed with smaller uninfected cells and the peripheral nodule cortex and parenchyma (Xiao et al., 2014). The latter contains the nodule vascular bundles. Due to the persistent activity of the meristem, a developmental gradient with distinct zones is formed along the longitudinal axis of the nodule (Hadri et al., 1998). Adjacent to the meristem is the infection zone, where cells, added by the meristem to the central tissue, can be penetrated by infection threads, from which rhizobia are released. Bacterial release requires the formation of unwalled droplets at the infection threads (Brewin, 2004). The released bacteria are surrounded by a host-derived membrane, by which organelle-like structures, called symbiosomes, are formed (Roth and Stacey, 1989). This requires the specific exocytotic vesicle-associated membrane proteins (VAMP721d&e) of the host plant (Ivanov et al., 2012). The release process occurs in about two cell layers adjacent to the meristem and in these cell layers the two NF receptors NFP and LYK3 accumulate (Moling et al., 2014). Both *in situ* hybridization and laser capture transcriptome data showed that nodulation (*nod*) genes, responsible for NF biosynthesis, are expressed in the apex of the nodule (Schlaman et al., 1991; Roux et al., 2014). Further, by a genetic approach it was shown that this expression occurs when rhizobia are still in the infection thread. Once released, bacteria no longer express *nod* genes (Marie et al., 1994). So, in the nodule, NF signaling only occurs at the apex, but it is not well known which genes are activated by NF signaling, and neither it is known which of these genes are involved in the release process.

Upon release rhizobia divide and gradually differentiate. In Medicago, bacterial differentiation involves endoreduplication, by which they become elongated. This is controlled by a number of nodule-specific cysteine-rich peptides (NCRs), which are produced by the host and are delivered to the rhizobia (Mergaert et al., 2003; Van de Velde et al., 2010; Alunni and Gourion, 2016). The infected host cells also undergo endoreduplication and become enlarged (Vinardell et al., 2013). The infection zone is followed by the fixation zone, where rhizobia fully differentiate and start to fix nitrogen.

The importance of NF signaling in nodules was shown by knockdown of the NF receptor gene *MtNFP,* which caused a block of bacterial release (Moling et al., 2014). It is further supported by knock down of *MtDMI2*, encoding a component of the NF signaling cascade, which resulted in a similar nodule phenotype (Limpens et al., 2005). Furthermore, the Medicago TE7 mutant, a weak allele of *IPD3* (orthologue of CYCLOPS), can form nodules, in which rhizobial release is completely blocked (Benaben et al., 1995a; Horváth et al., 2011; Ovchinnikova et al., 2011).

Although TE7 represents a weak IPD3 allele, NF signaling is sufficiently affected to block rhizobial release completely. This makes TE7 suitable to study NF signaling controlled gene expression in the nodule. However, after release of rhizobia from infection threads additional signaling between rhizobia and host might occur. This could result in additional changes in gene expression that would also be affected in TE7 nodules. Therefore, nodules in which bacterial release is blocked, but NF signaling is not affected can help to identify release depending transcriptional changes. Such nodules are formed on *VAMP721d&e RNAi* transgenic roots. Silencing these v-SNARE encoding genes, which are involved in a symbiotic exocytosis pathway, hindered the formation of unwalled droplets at the infection threads, and therefore blocking rhizobial release(Ivanov et al., 2012). NF signaling is likely not affected in these nodules. Therefore, comparing the transcriptomes of the nodule apical region where NF signaling and release occur in these mutant backgrounds, may allow the identification of genes regulated by NF signaling as well as the genes differentially expressed upon rhizobial release. Such genes can be directly regulated by NF signaling and indirectly regulated upon rhizobial release, or only regulated indirectly by signaling upon rhizobial release.

Here, we identified genes that are controlled by NF signaling (i.e. IPD3 dependent) in the nodule apex, as well as genes whose expression in addition depends on rhizobial release. This was done by analyzing the transcriptome of TE7, *VAMP721d&e* RNAi, and WT nodule apices obtained via laser microdissection. The identified set of NF-controlled genes in the nodule apex was markedly different from that regulated by NF signaling in the root. Further, we showed that NF signaling controlled rhizobial release depends on NIN and knock down of *NIN* is sufficient to block rhizobial release. This showed that NIN has a function in controlling rhizobial release, in addition to its multiple functions in the nodulation process previously identified.

## Materials and Methods

### Plant material and growth conditions

Medicago (*Medicago truncatula*) ecotype Jemalong 5 (J5) and A17, and TE7 mutant (Benaben et al., 1995) were used in this study. Plants were grown in a growth chamber at 20°C with 16h/8h day/night regime. Seed sterilization, germination and *Agrobacterium rhizogenes* msu440 mediated hairy root transformation was performed according to Limpens *et al*. (2004). Medicago plants were grown in perlite saturated with no nitrate (0 mM) containing Farhaeus (Fa) medium (Catoira *et al.,* 2000) for NIN RNAi knockdown experiment or low nitrate (0.25 mM Ca(NO3)_2_) for containing Farhaeus (Fa) medium for TE7 complementation experiment at 21°C and a 16 h : 8 h, light : dark regime. After 1 week of growth, plants were inoculated with *Sinorhizobium meliloti* 2011 (OD600 = 0.1, 2 ml per plant).

### Constructs

DNA fragments were PCR-amplified from *M. truncatula* A17 nodule genomic DNA or cDNA by using Phusion High-Fidelity DNA Polymerase (Finnzymes). The constructs were created by Gateway technology (Invitrogen). For knock down exocytosis pathway the *p35SCaMV:VAMP721d&e-RNAi* construct was generated in (Ivanov et al., 2012). For *pEnod12:NIN-RNAi* construct, the *Enod12* promoter was generated in (Limpens et al., 2005). The *NIN* fragment was cloned into pENTR-D-TOPO (Invitrogen) using NINRNAi-F (<u>CACC</u>GGAGAAAGTCCGGGGACAAG) and NINRNAi-R (CAAGCAGAGGGTGGGGATTT) primers. Then, it was recombined into the modified Gateway pK7GWIWG2(II)-*Q10::DsRED* binary vector. For p*IPD3:NIN* construct, the *NIN* coding sequence was cloned into pENTR-D-TOPO using NINcds-F (<u>CACC</u>ATGGAATATGGTGGTGGGTTAGTGG) and NINcds-R (GGAGGATGGACTGCTGCTGCTG) primers. The *IPD3* promoter (1,048 bp) in pENTR-p4p1r was generated in(Ovchinnikova et al., 2011). Next, a multisite GATEWAY reaction with LR Clonase II plus (Invitrogen) was performed using pENTR4-1-IPD3p, pENTR-D-TOPO-NIN, pENTR-p2rp3-Stop-T35S, and pKGW-RR-MGW (a modified multisite GATEWAY-compatible binary vector containing AtUbiquitin10p:DsRED1 to select transgenic roots) to generate *pIPD3:NIN*. The *pEnod12:GUS* construct was generated in (Li et al., 2021).

### Microscopy

Nodule formed after two or three weeks of growth in perlite was embedded in Technovit 7100 (Heraeus Kulzer), sectioning and staining were performed according to Xiao *et al*. (2014). Sections were analyzed with DM5500B microscope equipped with a DFC425C camera (Leica). Bright-field images were taken under a stereo macroscope (M165 FC, Leica). For electron microscopy, the tissue was fixed in 4% paraformaldehyde with 3% glutaraldehyde in 50 mM phosphate buffer, pH 7.4, postfixed with 1% OsO4, embedded in LR white resin according to the supplier’s recommendations and polymerized at 60 degree. Thin sections (60 nm) were cut using a Leica Ultracut microtome. The sections were contrasted with 2% aqueous uranyl acetate and lead citrate and examined using a JEOL JEM 2100 transmission electron microscope equipped with a Gatan US4000 4K × 4K camera.

### Laser microdissection

Nodules formed on J5, TE7 and *VAMP721d&e RNAi* were collected two weeks after inoculation with rhizobia. Further fixation, dehydration, embedding, section and laser microdissection procedure was performed according to (Zeng et al., 2018). Leica LMD7000 laser microdissection microscope was used for laser capture. About 2000 cells were collected for each replicate.

### RNA isolation and quality check

RNA was isolated using the Qiagen RNeasy Micro kit according to manufacturer’s protocol. The quality of isolated RNA samples was checked using both Agilent 2100 Bioanalyzer and qPCR. For quality check by qPCR, 2 µL of isolated RNA sample was reverse transcribed using iScript cDNA Synthesis kit (Bio-Rad) and pre-amplified using Bio-Rad Sso Advanced PreAmp Supermix. *NIN*, *NF-YA1*, *VAMP721d* and *VAMP721e* were used as marker genes checked by qPCR. RNA samples from the best three replicates of each cell type were amplified using the SMART-Seq v4 Ultra Low Input RNA Kit (Clontech, http://www.clontech.com) according to manufacturer’s instructions. The number of amplification cycles (11–15 cycles) for each sample was adjusted based on the amount of RNA determined by bioanalyzer and qPCR analyses. Real-time qPCR was performed in 10 µl reactions using SYBR Green Supermix (Bio-Rad) and a CFX real-time system (Bio-Rad). Gene expression was normalized using *ACTIN2* as a reference gene.

### Library preparation and sequencing

Sequencing libraries were made using ThruPLEX DNA-seq Kit (Rubicon genomics, http://rubicongenomics.com/) and sequenced on an Illumina Novaseq 6000 platform (paired-end) at the Next-Generation Sequencing (NGS) facility of Bioscience, Wageningen University & Research.

### RNA-seq analyses

About 4 Gb cleaned reads were obtained for each sequenced sample. Sequence reads were mapped to the *M. truncatula* genome v.5.0 (Pecrix *et al.,* 2018) using the CLC genomics workbench 10.0.1 (Qiagen). Length fraction and similarity fraction were set to 0.9 during mapping and only uniquely mapped reads were considered in the analysis. All other parameters were set as default. Differential expression analyses were performed using the CLC genomic workbench 10.0.1. The criteria of fold change > 2, false discovery rate (FDR) P-value < 0.05 and average TPM of J5, TE7 and *VAMP721d&e RNAi* > 3 were used to identify the differentially expressed genes in TE7 nodule apices. Because studies on *VAMP721d&e RNAi* nodules relied on hairy root transformation, which have a higher degree of variation between the three replicates, we use the criteria of fold change > 2, FDR P-value < 0.2 and average TPM of J5, TE7 and *VAMP721d&e RNAi* > 3 to identify the differentially expressed genes in *VAMP721d&e RNAi* nodule apices.

## Results and Discussion

### Identification of genes regulated by NF signaling in nodules

NF receptors accumulate and form heterodimers in Medicago nodules, but only in two cell layers adjacent to the apical meristem (Moling et al., 2014). To identify genes that are controlled by NF signaling in nodules, we made use of laser microdissection to specifically enrich for these cell layers (Fig. S1). Rhizobial release occurs in the same cell layers, therefore, to identify NF signaling controlled genes and the genes that are regulated, in addition, upon release of rhizobia, we made use of three Medicago lines: WT (J5) (release+/NF signaling+), *p35SCaMV:VAMP721d&e RNAi* (release-/NF signaling+) and TE7 (release-/NF signaling-) (Fig. 1a) (Ovchinnikova et al., 2011; Ivanov et al., 2012). RNA was extracted from laser captured nodule apices (Fig. S1) of these Medicago lines (three biological replicates for each line) and it was amplified for library preparation and subsequent RNA-seq.

**Fig. 1.**
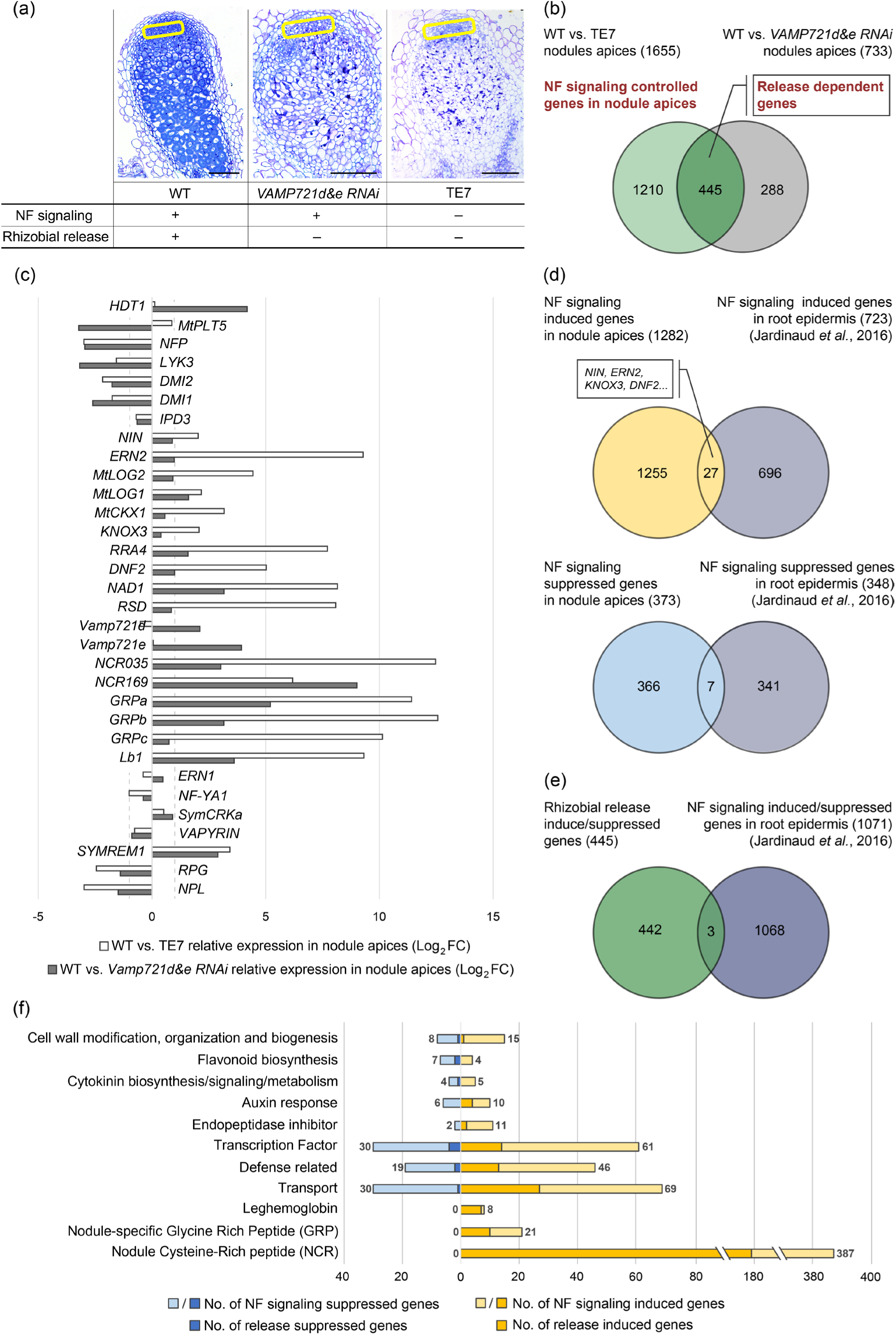
Genes controlled by NF signaling in root nodule apices and a comparison with genes regulated by NFs in root epidermis. (a) Characteristics of the mutant nodules used for RNA-seq. Semi-thin sections showing that rhizobial release is blocked in TE7 and *VAMP721d&e RNAi* nodules. The sections were stained with toluidine blue. The yellow rectangle outlines indicate the laser-captured areas. Scale bars: 150 μm. (b) A Venn diagram showing the number of genes differentially expressed in TE7 nodules apices (1655 genes), and the number of genes differentially expressed in *VAMP721d&e RNAi* nodules apices (733 genes) (Table S1a-1c). The genes differentially expressed in TE7 nodule apices in comparison to WT are most likely NF signaling controlled genes (Table S1a). The 445 genes, which are differentially expressed in both TE7 and *VAMP721d&e RNAi* nodule apices in comparison to WT, are likely genes whose expression depends on release (Table S1b). (c) Analysis of representative genes from the transcriptome data (Table S1d). (d) Venn diagrams showing that the overlap between NF signaling regulated genes in the root epidermis and NF signaling controlled genes in nodules is small (Table S2a and 2b). The upper panel represents induced genes; the lower panel represents suppressed genes. (e) A Venn diagram showing that 442 out of 445 putative release-dependent genes were not shared with genes that have an altered expression level in the root epidermis in response to NF application (Table S2d). (f) NF signaling regulated gene families and metabolism pathways in Medicago nodule apices. The classification of each gene in the families and metabolism pathways is based on the protein function indicated in the Mt 5.0 annotation or gene ontology terms (Table S3).

Genes controlled by NF signaling are most likely differentially expressed in TE7 nodule apices in comparison to WT nodule apices. In total, we identified 1655 of such differentially regulated genes (Fig. 1b, Table S1a). Among these genes, 373 were upregulated and 1282 were down regulated in TE7. Since release of bacteria from the infection threads is blocked in the TE7 nodules, genes that are regulated upon release would be among the differentially expressed genes. These release dependent genes are directly or indirectly regulated by NF signaling. To determine which NF signaling controlled genes depend on rhizobial release, we made use of *VAMP721d&e RNAi* nodules. Rhizobial release is blocked in *VAMP721d&e RNAi* nodules due to reduced activity of a symbiotic exocytosis pathway (Ivanov et al., 2012) and likely not due to impaired NF signaling. Therefore, the genes that are differentially expressed in both TE7 and *VAMP721d&e RNAi* nodule apices in comparison to WT are likely release dependent genes. By this comparison, we identified 445 genes regulated upon rhizobial release (Fig. 1b, Table S1b). Because studies on *VAMP721d&e RNAi* nodules relied on hairy root transformation, in which each root is the result of a different transformation event, some release might have taken place in some nodules. So some release dependent genes might be missed in this comparison. It is noteworthy that 288 genes are differentially expressed in *VAMP721d&e RNAi,* but not in TE7 nodule apices in comparison to WT (Fig. 1b, Table S1c). These include for example HDT1 and PLT5 (Fig. 1c). There can be several reasons why they are only differentially expressed in *VAMP721d&e RNAi* nodule apices. For example, when genes are induced by NF signaling, but suppressed upon release, they can be differentially expressed in *VAMP721d&e RNAi* but not in TE7. Alternatively, it is possible that VAMP721d&e are, in addition to release, involved in another process. Since these possibilities are indistinguishable with the current data, we will not focus on these genes.

Among the genes regulated upon rhizobial release we observed the gene encoding NF receptor NFP, which is suppressed upon rhizobial release (i.e. higher expression in TE7 and *VAMP721d&e RNAi* compared to WT nodule apices) (Fig. 1c, Table S1b). Similar suppression also holds for genes encoding NF receptor LYK3 and the downstream NF signaling component DMI2 and DMI1, although less significant (Fig. 1c, Table S1d). It shows that rhizobial release correlates with a negative regulation of NF signaling components, and this is consistent with the narrow region in which NF receptor genes and *DMI2* are expressed in the nodule apex (Limpens et al., 2005; Moling et al., 2014). We will describe the other representative genes shown in Fig. 1c after the comparison of NF signaling controlled genes in nodule and root epidermis.

### NF signaling controls a markedly different set of genes in nodules in comparison to roots

NF signaling initially occurs at the onset of nodulation when roots are inoculated with rhizobia (or treated with NFs), and it has been well studied which genes are differentially expressed at this stage. We decided to compare the IPD3-dependent NF signaling controlled genes in nodule apices with NF signaling regulated genes in roots. Previously, it has been reported that 1071 genes were differentially expressed in the root epidermis in response to 4 or/and 24 hours of NF treatment (Jardinaud et al., 2016). We compared these 1071 genes with the 1655 genes whose expression is regulated by NF signaling in nodule apices. The latter only shared 27 genes that were induced and 7 genes that were suppressed in the root epidermis upon NF application (Fig. 1d, Table S2a and 2b). To test to what extent this limited overlap is caused by other bacterial signals besides NFs, we repeated the analysis using microarray data of Medicago root hairs 1, 3, and 5 days after inoculation with WT rhizobium or a mutant unable to produce NFs (Breakspear et al., 2014). This comparison showed similar results. Among the 485 genes induced in roots by NF producing rhizobia, only 9 genes overlapped with the genes induced by NF signaling in nodule apices and none of the genes suppressed in roots was suppressed in nodules (Fig. S2, Table S2c). Together, this suggests that the set of genes controlled by NF signaling in nodules is markedly different from that in roots.

Genes that are induced by NF signaling in both root epidermis and nodule apices include *NIN*, *ERN2*, genes involved in cytokinin biosynthesis/signaling such as *KNOX3* and A-type *RR4*, genes involved in defense suppression such as *DNF2*, auxin response genes, and genes involved in flavonoid biosynthesis (Fig. 1c and 1f, Table S2a-2c). These genes appear to depend less on cellular/developmental conditions, maybe because they are primary targets of NF signaling (Singh et al., 2014a; Soyano et al., 2014; Liu et al., 2019). Although for some genes, this remains to be demonstrated.

It is noticeable that there were many genes that were regulated by NF signaling in roots, but whose expression is not affected in the TE7 nodule apex. Those are for example, *ERN1*, *NF-YA1*, *SymCRK*, and *VAPYRIN* (Fig. 1c, Table S1d). Furthermore, there were some genes induced by NF signaling in roots but showed higher expression in TE7 nodule apices compared to WT, such as *RPG* and *NPL* (Fig. 1c, Table S1a). There can be several reasons to explain these. First, the expression of these genes might fully depend on NF signaling in the root epidermis that sets in motion the nodule developmental program. In *Medicago sativa* (alfalfa) and *Vicia sativa* (vetch) it has been shown that application of NF can induce formation of nodules and some nodule specific genes are expressed in these nodules (Truchet et al., 1989; Vijn et al., 1995). As NFs are immobile molecules (Goedhart et al., 2000), transcriptional changes in these nodules must be the result of initial NF signaling in the root epidermis. So, NF signaling at the epidermis sets in motion a developmental process, resulting in nodule formation in which some nodule specific genes are expressed. Therefore, the expression of genes like *ERN1*, *NF-YA1*, *SymCRK*, and *VAPYRIN* in TE7 nodules might be the result of the intrinsic nodule developmental program that is initiated by NF signaling at the epidermis. Alternatively, because TE7 is a weak allele and it still has its functional redundant *IPD3-like* gene, a low level of NF signaling might be sufficient to trigger their expression. Furthermore, in WT nodules their induction by a higher level of NF signaling might be followed by negative feedback induced upon rhizobial release. In this case their expression level can be equal in TE7 and WT, or even higher in TE7 nodules.

A major difference between NF signaling in roots and nodules is its involvement in triggering release of bacteria from the infection threads to form symbiosomes. Such release only occurs in the nodule and not in root cells. This difference was reflected in the observation that the vast majority of release dependent genes (442 out of 445) were not shared with NF regulated genes in the root epidermis (Fig. 1e, Table S2d). Therefore, occurrence of rhizobial release in the nodule is a major contribution to the large difference. The remaining difference between nodule and root might be caused by differences in cellular/developmental conditions. It might be that such genes require different levels of NF signaling or that the cellular environment, for example through combinatorial action of different transcription factors and or epigenetic differences, determines which genes are regulated.

To understand what might be the biological reason for the different sets of genes regulated by NF signaling in nodules and roots, we looked into the function of genes regulated by NF signaling. These include for example 387 *NCR* genes and 21 *GRP* genes, which were induced by NF signaling directly or indirectly in the nodule apices (Fig. 1f, Table S3). Several of these genes have been shown to be involved in rhizobial differentiation into elongated nitrogen fixing bacteroids (Kevei et al., 2002; Pan and Wang, 2017). Further, NF signaling induces eight genes encoding Leghemoglobins, which facilitate oxygen delivery to the symbiosomes and are essential for nitrogen fixation (Ott et al., 2005; Wang et al., 2019). Expression of seven of these genes was release dependent (Fig. 1f, Table S3). Other examples of NF signaling induced genes are genes required for bacterial intracellular accommodation, such as the defense suppressing genes *DNF2*, *NAD1* and RSD (Sinharoy et al., 2013; Bourcy et al., 2013; Wang et al., 2016) (Fig. 1c, Table S1a). It is noticeable that although these defense suppressing genes were highly induced in WT nodule apices by NF signaling, many defense response genes were still higher expressed in WT in comparison to TE7 nodule apices (Fig. 1f, Table S3). The released rhizobia in WT nodules are likely responsible for induction of these defense response genes. 13 out of 46 defense response genes were release dependently induced. Some of the release dependent defense response genes might be missed because release occurred in some *VAMP721d&e RNAi* nodules. Rhizobial release requires cell wall modifications. Consistent with this, we identified 23 cell wall modification, organization and biosynthesis genes were regulated by NF signaling in nodules (Fig. 1f, Table S3). This might include genes essential for rhizobial release. Although NF signaling in the nodule only occurred in one to two cell layers adjacent to the nodule meristem, its effect on transcriptional changes likely last for subsequent nodule developmental stages. NF signaling controls the expression of 91 transcription factor encoding genes in nodule apices. These transcription factors likely will induce further transcriptional changes. So, NF signaling in the nodule regulates many genes that are required for processes that occur in nodules, but not in roots. The expression of some of these genes depends on rhizobial release.

Here we showed that, different from NF signaling controlled genes in roots, NF signaling in nodules control a subset of genes that are required for rhizobial differentiation and accommodation in nodules. Furthermore, we showed that rhizobial release in the nodule contributes to the induction of many nodule-specific genes. This suggests that additional signaling events between plant and bacteria are occurring after release or that a developmental switch, which requires NF signaling, also depends on the release of the bacteria from the infection threads. Our study is based on Medicago, which forms indeterminate nodules(Hirsch, 1992). However, many legume species such as Lotus, form determinate nodules, in which the meristem is only transiently present. In these nodules, a developmental gradient is not formed along the longitudinal axis (Hirsch, 1992). It will be interesting to see whether NF receptors also accumulate in these determinate nodules and to what extent they control gene expression in this nodule type.

### Nod factor signaling induced rhizobial release depends on NIN

One of the genes that was significantly less induced in the TE7 nodule apex, but not affected in the *VAMP721d/e RNAi* nodules, was the key NF signaling gene *NIN*. As a primary target of CYCLOPS (orthologue of MtIPD3) (Singh et al., 2014b), the induction of *NIN* at a too low level at the nodule apex might have caused the block of rhizobial release in TE7 (*ipd3* mutant). To test this, we introduced *NIN,* driven by the functional *IPD3* promoter (*pIPD3:NIN*) (Ovchinnikova et al., 2011), into TE7 by hairy root transformation. Three weeks after inoculation with rhizobia, 44 out of 81 nodules formed on roots transformed with *pIPD3:NIN* showed release of rhizobia (Fig. 2a-2b). Such release was not observed in any of the nodules formed on the TE7 roots transformed with the empty vector (n=67) (Fig. 2c-2d). Quantitative RT-PCR showed that the *NIN* transcript level was about four times higher in nodules formed on TE7 roots transformed with *pIPD3:NIN* compared with the nodules formed on roots transformed with empty vector (Fig. S3a). A similar rhizobial release was observed when we introduced *pIPD3:NIN* into *ipd3-2,* a strong *ipd3* allele (Fig. S3d-3i). These results suggest that the block of bacterial release in TE7 is the result of a too low expression level of *NIN*.

**Fig. 2.**
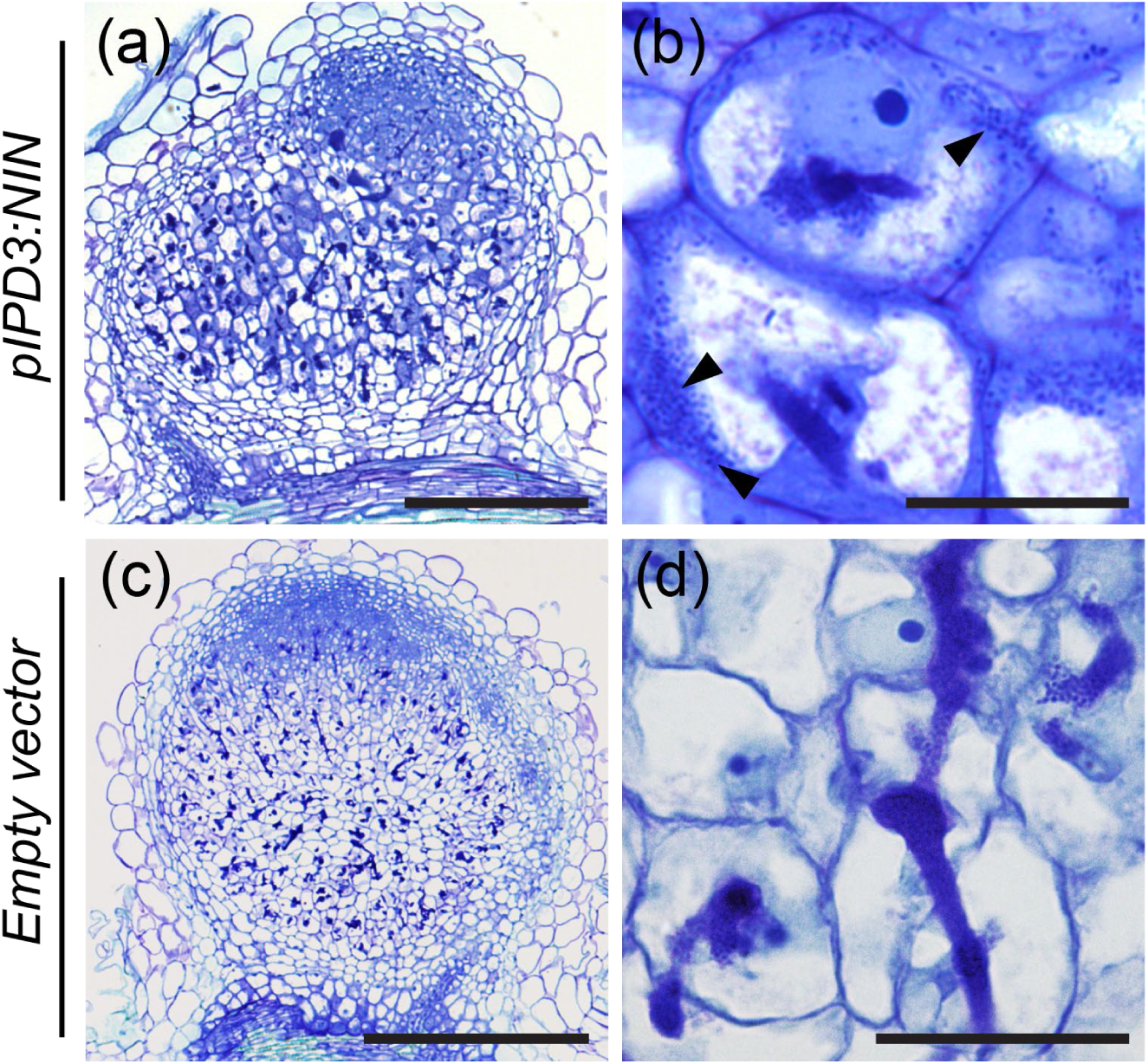
*pIPD3:NIN* complements the block of bacterial release phenotype in the TE7 mutant. (a) A nodule formed on TE7 roots transformed with *pIPD3:NIN* at 3 weeks post inoculation (wpi). (b) Magnification of infected cells of this nodule, showing bacteria (arrowheads) released into plant cells. (c) A nodule formed on TE7 roots transformed with Empty vector at 3 wpi. (d) Magnification of infected cells of this nodule, showing no bacterial release. Semi-thin nodule sections stained with toluidine blue. Scale bars: (a, c) 300 μm; (b, d) 30 μm.

Further, we tested whether a reduction of *NIN* expression level is sufficient to cause a block of release. To test this, we used a conditional knockdown (RNAi) approach. By hairy root transformation we introduced a *NIN* RNAi construct, driven by the nodule specific *Enod12* promoter, into WT (A17) roots. This *Enod12* promoter is expressed in nodule primordia (Li et al., 2021) and in the apical part of the nodule (Fig. S3b), including the region where rhizobial release occurs. Two weeks after inoculation, only a few nodules were formed on *pEnod12:NIN RNAi* transgenic roots. This is consistent with NIN being essential during nodule initiation. These nodules were small and white, indicating that their development was hampered (Fig. 3a). Quantitative RT-PCR showed that the *NIN* transcript level was reduced to 27% in *NIN* RNAi nodules compared with the control nodules (Fig. S3c). Semi-thin sections of these nodules showed that in 17 out of 38 nodules, release was blocked (Fig. 3c and 3e). In contrast, none of the analysed nodules (n=39) formed on roots transformed with empty vector showed such phenotype (Fig. 3d and 3f). Block of rhizobial release in *NIN* RNAi nodules indicated that the formation of an unwalled droplet, a structure required for rhizobial release, is affected. To check this, we further studied the *NIN* RNAi nodules by electron microscopy. We compared *NIN* RNAi nodules, where rhizobial release is blocked, with nodules formed on control roots. In the latter, unwalled droplet structures were formed (Fig. 3h and 3j). In the *NIN* RNAi nodules, infection threads have structures that resemble infection droplets. However, these droplet-like structures were surrounded by a wall (Fig. 3g and 3i), which prevented the contact between the bacteria and the host membrane and thereby hampered the pinching off of symbiosomes.

**Fig. 3.**
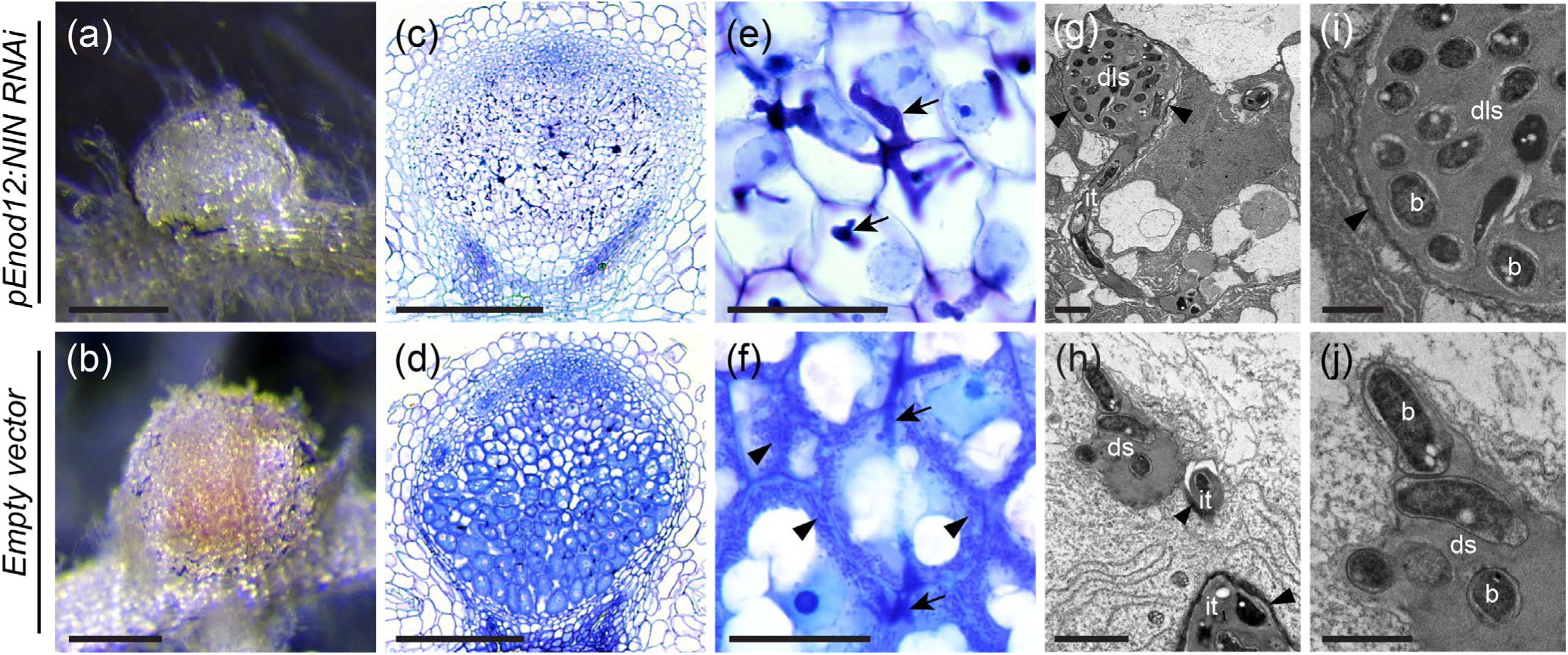
knockdown of *NIN* in the nodule apex blocks rhizobial release. (a) and (b) Pink nodules were formed on Medicago WT (A17) roots transformed with empty vector (b), whereas nodules formed on *pEnod12:NIN RNAi* transgenic roots were small and white (a). (c) to (f) Semi-thin nodule sections stained with toluidine blue. [(c) and its magnified image (e)] 17 out of 38 *pEnod12:NIN RNAi* nodules have many infection threads (arrow) formed, but the release of rhizobia into the plant cell were hampered, as no symbiosomes were observed In contrast, in nodules formed on roots transformed with empty vector bacteria were released from infection threads (arrow) into plant cells, and symbiosomes (arrowhead) were formed [(d) and its magnified image (f)]. (g) and its maginfied image (i) Electron microscopy images of an infection thread in a *pEnod12:NIN RNAi* nodule. At the tip of the infection thread (it) a droplet-like structure (dls) was formed. This droplet-like structure is filled with bacteria (b) and surrounded by a wall (arrowhead). (h) and its magnified image (j) Electron microscopy images of infection threads (it) in a control nodule. Infection thread wall indicated by arrowheads. Bacteria (b) were released from an unwalled droplet structure (ds). Scale bars: (a, b) 2mm; (c, d) 300 μm; (e, f) 30 μm; (g, h) 2 μm; (i, j) 1 μm.

These results showed that NF signaling controlled rhizobial release depends on NIN. This most likely occurs in a cell autonomous manner as *NIN* is expressed in the cell layers where release and NF signaling occur. However, the expression of *NIN* is not restricted to the two cell layers where NF signaling takes place. *NIN* is expressed in the complete infection zone and its expression level gradually increases till the transition to the fixation zone when it suddenly drops to a very low level (Liu et al., 2021). NIN is not only involved in rhizobial release, but also in differentiation of symbiosomes, suppression of defense and premature senescence and induction of the expression of *leghemoglobin* genes (Jiang et al., 2021; Liu et al., 2021). The expression pattern of *NIN* and its multiple functions in root nodules illustrates how NF signaling sets in motion a developmental process by which functional nodule cells are formed. It starts with NF signaling in a very narrow region next to the meristem. This results in the induction of rhizobial release and expression of many genes including *NIN* in a cell autonomous manner. When cells leave this narrow region NF signaling is suppressed, but genes like *NIN* can maintain (and even increase) their expression. The nodule cells and symbiosomes gradually differentiate by which the functional part of the nodule, the fixation zone, is formed. So the genes that we classified as NF signaling induced genes are in principle induced in a cell autonomous manner. These will result in further transcriptional changes at subsequent nodule developmental stages.

## Acknowledgements

This research was supported by the European Research Council (ERC-2011-AdG-294790), and the China Scholarship Council (201506300062 to JL).

## Author contributions

J.L., E.L., R.H. and T.B. designed the research. J.L., D.S., E.L., E.E.F., R.H., H.Li., H.C.B., E.S., O.K. performed experiments. J.L., E.L., E.E.F T.Z., O.K. and T.B. analysed data. J.L. and T.B. wrote the manuscript.

## Data availability

Sequence data from this article have been submitted to Sequence Read Archive (SRA). The accession numbers are SRR25037585; SRR25037584; SRR25037583; SRR25037582; SRR25037581; SRR25037580; SRR25037579; SRR25037578; SRR25037577; SRR25037576; SRR25037574; SRR25037575; SRR25037573; SRR25037572; SRR25037571; SRR25037570; SRR25037569; SRR25037568

## Supporting Information

**Fig. S1** Laser capture microdissection of regions adjacent to the nodule meristem.

**Fig. S2** Venn diagrams showing a small overlap between NF signaling controlled genes in nodules and those in root epidermis.

**Fig. S3** Supporting information for Nod factor signaling induced rhizobial release depends on NIN.

**Table S1** Identified NF signaling regulated genes in nodules. (separate Excel file)

**Table S2** Overlapped genes between NF signaling regulated genes in root epidermis and in nodules. (separate Excel file)

**Table S3** NF signaling regulated gene families and metabolism pathways in Medicago nodule apices. (separate Excel file)

