## Supplementary material for "Nod factor signaling controlled genes in *Medicago truncatula* nodules": Fig. S1-3

Article acceptance date: Click here to enter a date.

The following Supporting Information is available for this article:

**Fig. S1** Laser capture microdissection of regions adjacent to the nodule meristem.

**Fig. S2** Venn diagrams showing a small overlap between NF signaling controlled genes in nodules and those in root epidermis.

**Fig. S3** Supporting information for Nod factor signaling induced rhizobial release depends on NIN.

**Fig. S1** Laser capture microdissection of regions adjacent to the nodule meristem. Representative nodule samples from WT (J5) (a and b), TE7 (c, d and f), and *VAMP721d&e RNAi* (e)*,* before (a, c and e), and after (b and d) laser-microdissection. (f) Collected samples of the cell layers adjacent to the apical meristem, including few neighbouring cell layers.

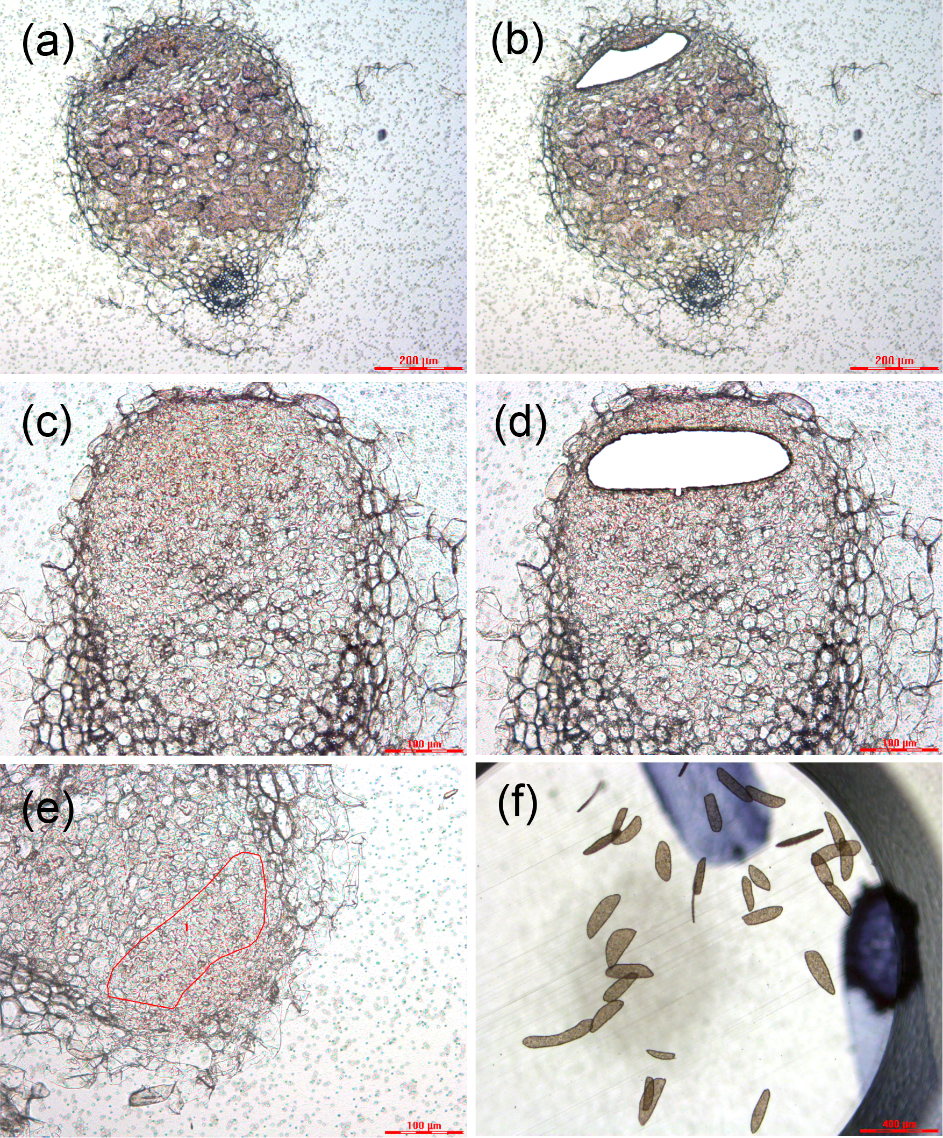

**Fig. S2** Venn diagrams showing a small overlap between NF signaling controlled genes in nodules and those in root epidermis.

Upper panel: comparison of NF signaling induced genes. Lower panel: comparison of NF signaling suppressed genes. Overlapped genes shown in Table S2c.

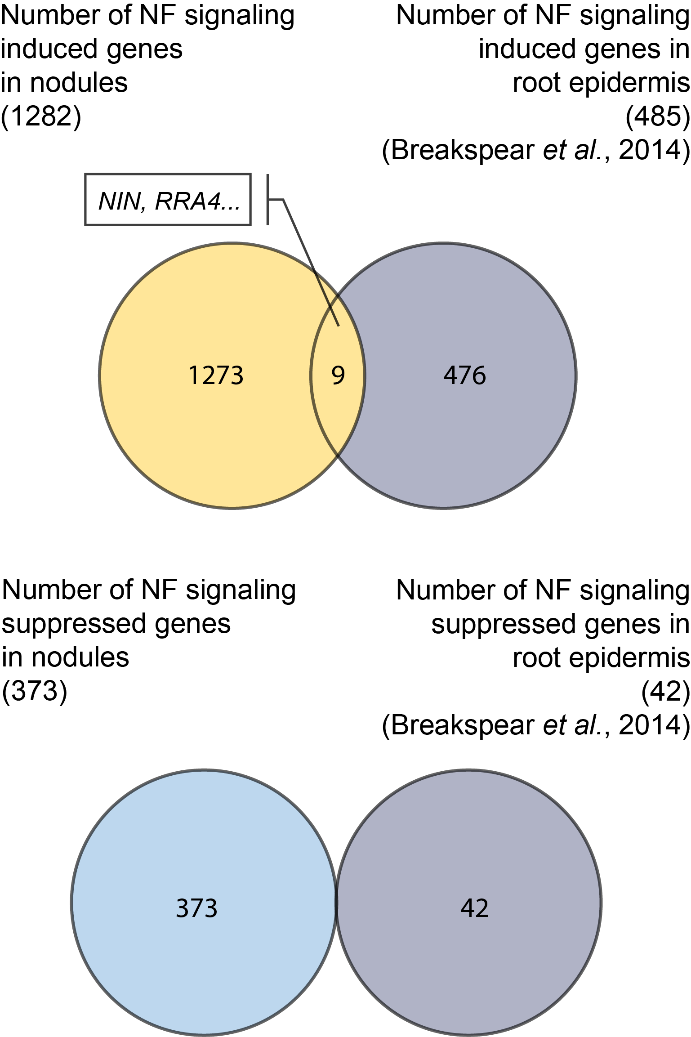

**Fig. S3** Supporting information for Nod factor signaling induced rhizobial release depends on NIN. (a) *NIN* transcript level increased in nodules transformed with *pIPD3:NIN*. Quantitative reverse transcription polymerase chain reaction (qRT-PCR) showed that the *NIN* transcript level was increased about four times in nodules formed on TE7 roots transformed with *pIPD3:NIN* compared with nodules formed on TE7 roots transformed with empty vector. Data are means $\pm$SD of three biological replicates.

(b) *pEnod12:GUS* expression pattern in A17 nodules. GUS activity was observed in the apical part of the nodule, including the adjacent cell layers of meristem and infection zone, where rhizobial release occurs. M, meristem; IF, infection zone; FX, fixation zone. Scale bar: 100 $\mu$m.

(c) *NIN* transcript level is reduced in nodules transformed with *pEnod12:NIN RNAi.* Quantitative reverse transcription polymerase chain reaction (qRT-PCR) showed that the *NIN* transcript level was reduced to 27% in *NIN* RNAi nodules compared with the nodules transformed with empty vector. Data are means ± SD of three biological replicates.

(d-i) *pIPD3:NIN* complements the infection and release phenotype of the *ipd3-2* mutant.

(d) At 3 wpi, elongated nodules were formed on *ipd3-2* roots transformed with *pIPD3:NIN*.

(e) Infected nodules formed in on *ipd3-2* roots transformed with *pIPD3:NIN.* (f) Magnification of infected cells, showing bacteria (arrowheads) released into plant cells.

(g) At 3 wpi, only bumps were formed on *ipd3-2* roots transformed with empty vector. (h) Section of such structures revealed that it is not infected. This is further confirmed in the zoom up image (i).

Semi-thin nodule sections stained with toluidine blue (e), (f), (h) and (i). Scale bars: (d, g) 2 mm; (e, h) 200$\mu$m; (f, i) 20$\mu$m.

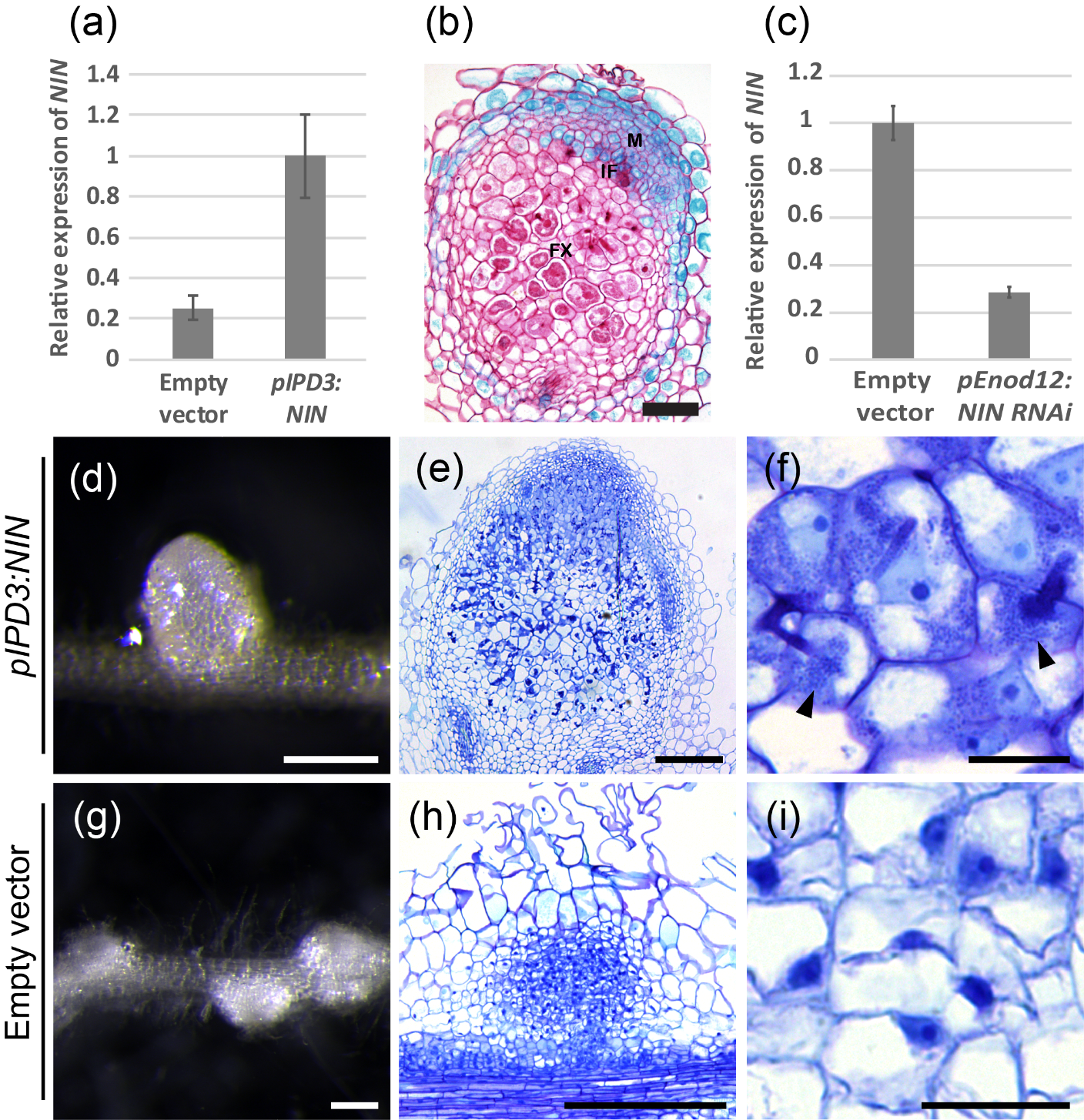
